# Evolutionary emergence and preservation of microproteins encoded by upstream ORFs

**DOI:** 10.64898/2026.03.05.709866

**Authors:** José Carlos Montañés, Chris Papadopoulos, Saif Al-Obaidi, Anikó Szegedi, William R. Blevins, Marc Talló-Parra, Juana Díez, Elena Hidalgo, M. Mar Albà

**Affiliations:** Evolutionary Genomics Group, Hospital del Mar Research Institute, Barcelona, Spain; Bioinformatics Knowledge Center, Howest University of Applied Sciences, Bruges, Belgium; Centro Nacional de Análisis Genómico (CNAG), Barcelona, Spain; Molecular Virology Group, Department of Medicine and Life Sciences, Universitat Pompeu Fabra (UPF), Barcelona, Spain; Oxidative Stress and Cell Cycle Group, Universitat Pompeu Fabra (UPF), Barcelona, Spain; Catalan Institution for Research and Advanced Studies (ICREA), Barcelona, Spain

## Abstract

The analysis of ribosome profiling (Ribo-Seq) data has provided evidence that many eukaryotic mRNAs contain translated upstream or downstream ORFs (uORFs/dORFs), but the biological significance of this translation activity remains, for the most part, unknown. One of the principal limitations has been the lack of Ribo-Seq data from several closely related species, precluding the identification of cases in which translation is phylogenetically conserved. Here, by combining Ribo-Seq data from 100 different experiments, we identify 2,332 translated uORFs and 1,008 translated dORFs in *S. cerevisiae*, which result in microproteins that tend to be highly hydrophobic or positively charged. To study their phylogenetic conservation, we have generated Nanopore direct RNA sequencing data, together with Ribo-Seq data, from six additional *Saccharomyces* species, spanning an evolutionary period of around 16 million years. We have identified 195 translated *S. cerevisiae* uORFs that are also translated in other *Saccharomyces* species; these uORFs are translated at levels comparable to the main coding sequence and display signatures of purifying selection at the level of the encoded microproteins. In contrast, dORFs are translated at very low levels and they are rarely conserved, suggesting much more limited microprotein functionalization. We have also discovered that uORF translation is associated with the formation of alternative transcript isoforms encompassing the region containing the uORFs but not the main protein coding sequence, implying that some microproteins can be produced independently of the main protein product. This work significantly advances our understanding of how initially pervasive uORF translation can result in new microproteins, providing many new candidates for further functional studies.

## INTRODUCTION

Recent studies have shown significant translation of many non-canonical open reading frames (ncORFs) outside annotated coding sequences (Ingolia et al. 2009; Blevins et al. 2021; Chen et al. 2020; Patraquim et al. 2020; Mudge et al. 2022; Chothani et al. 2022; Saghatelian and Couso 2015). The largest class is constituted by upstream ORFs (uORFs), located in the 5’untranslated region (5’UTR). Many mRNAs also contain translated downstream ORFs (dORFs), located in the 3’UTR, with overlapping upstream ORFs (ouORFs) and overlapping downstream ORFs (odORFs) occurring at lower frequencies. With a few exceptions, the functional consequences of the translation of ncORFs in UTRs, if any, are unknown.

The majority of the translated ncORFs, including uORFs, show limited sequence conservation across species (Sandmann et al. 2023); this implies a lack of function, i.e. that their translation is merely a by-product of other processes. However, a study of sequence variants that create or disrupt uORFs in humans suggested that some instances are indeed preserved by purifying selection (Whiffin et al. 2020). There are several known human disease which stem from mutations in uORFs (Lee et al. 2021). Additionally, research which compared intra-specific diversity *versus* inter-specific divergence of uORF sequences from different eukaryotic species have concluded that positive selection is likely to have shaped their evolution (Zhang et al. 2018, 2021).

Several cases have been described in which the translation of uORFs can inhibit the translation of the main coding sequence (CDS). One such example is the stress master regulator *GCN4*, whose translation is rapidly activated during stress due to changes in the occupancy of ribosomes at uORFs (Grant, Miller, and Hinnebusch 1995; Hinnebusch, Ivanov, and Sonenberg 2016; Hinnebusch 2005). However, the level of translation of uORFs and the main coding sequence in different conditions are generally positively correlated (Heesch et al. 2019; Patraquim et al. 2020; Moro et al. 2021), suggesting that the majority of the uORFs are probably not regulatory. Evidence has also accumulated in recent years that some microproteins encoded by uORFs can have a function *per se* (Andrews and Rothnagel 2014; Renz et al. 2020) - for example, the human microprotein MIEF1 regulates mitochondrial translation (Rathore et al. 2018). In addition, a recent study using CRISPR-Cas uORF deletion experiments in human cell lines detected that, in several cases, uORF deletion resulted in decreased cell growth (Chen et al. 2020).

In this study we use long read transcriptomics data and Ribo-Seq data from the unicellular eukaryotic model organism *Saccharomyces cerevisiae* and six other closely related species to shed new light into the process of emergence and functionalization of microproteins encoded by translated uORFs in mRNAs.

## RESULTS

### Reconstruction of full transcripts using Nanopore dRNA

Our first goal was to fully reconstruct the transcriptome of baker’s yeast (*Saccharomyces cerevisiae*), alongside six other *Saccharomyces* species, spanning the diversity of the genus (Figure 1a)(Wolfe 2006; Hittinger 2013; Peris et al. 2023). The most distant species was *S. uvarum*, which split from *S. cerevisiae* about 16 Million years ago (Marcet-Houben and Gabaldón 2015). The genome assemblies of the seven *Saccharomyces* species are complete, but the annotations are mostly limited to coding sequences. In order to obtain the 5’UTR and 3’UTR annotations for all mRNAs, we performed Nanopore direct RNA (dRNA) sequencing for all seven species (between 33-49 million reads)(Table S1). After polishing the dRNA reads with Illumina RNA-Seq data, we generated new transcriptome assemblies that included the 5’ and 3’UTR regions in addition to the coding sequence (CDS)(Figure 1b). These comprehensive annotations are available for all seven species as supplementary material.

**Figure 1.**
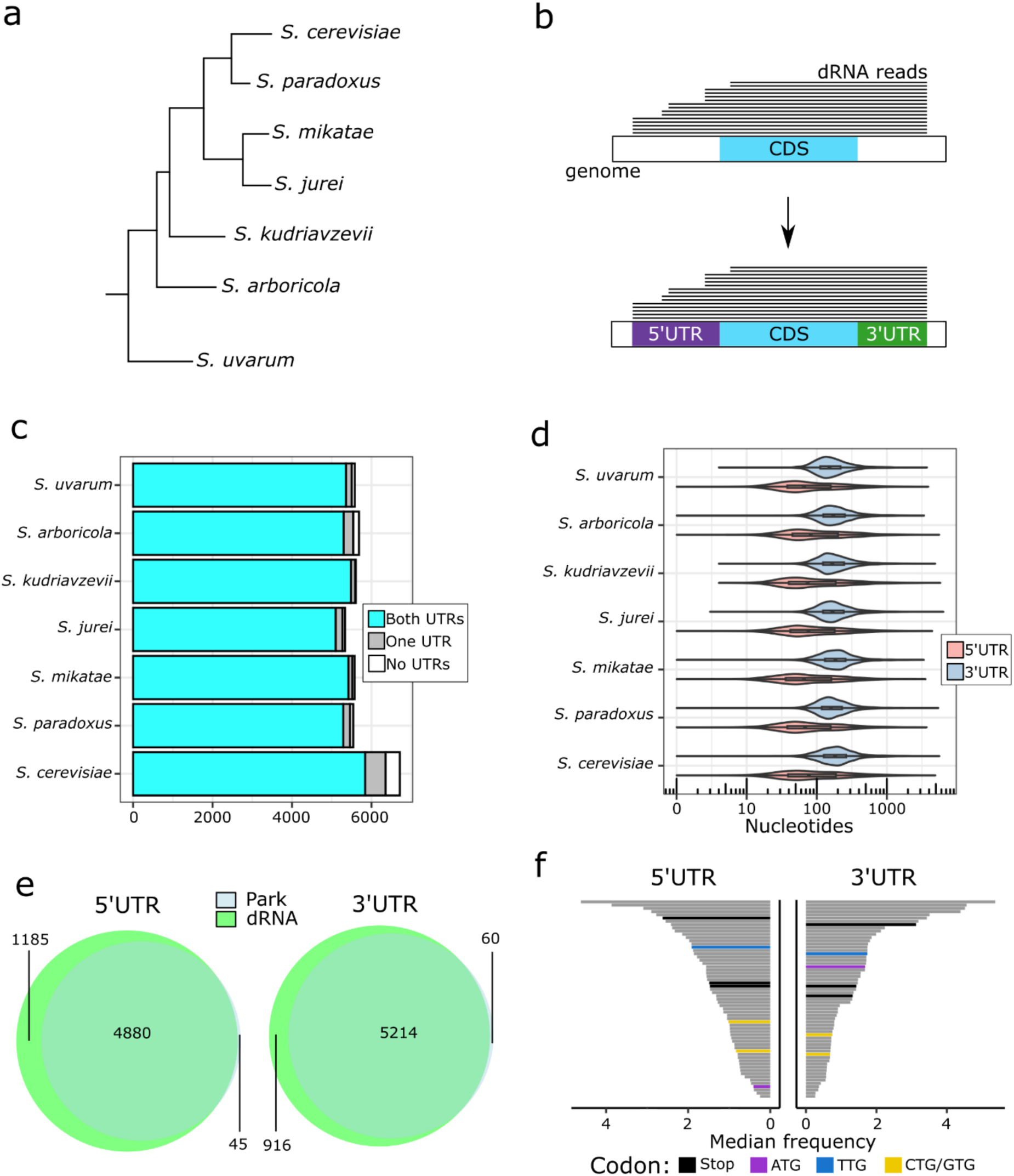
Annotation of 5’ and 3’UTRs using Nanopore dRNA. **(a)** Phylogenetic representation of the *Saccharomyces* species analyzed in this study - branch lengths are not representative of evolutionary distance. **(b)** Nanopore direct RNA (dRNA) reads were mapped to the reference genomes and used to generate the UTRs of transcribed genes. **(c)** Number of UTRs reconstructed for the different species (see also Table S2). **(d)** Size distribution of 3’ and 5’ UTRs in the different species (see Table S3 for medians and quartiles). **(e)** Number of genes in which we identified UTRs in *S. cerevisiae versus* those identified by Park, et al. 2014. **(f)** Median frequency of different triplets in the 5’ and 3’ UTR of *S. cerevisiae* genes. The triplet type is indicated by colored bars at the bottom of the figure. Gray bars represent the remainder of triplets.

Our methodology recovered 5’UTR and 3’UTR sequences for the vast majority of the mRNAs (Figure 1c)(Table S2). The 5’UTR and 3’UTR had a similar overall size distribution in the different species (Figure 1d, Table S3). For *Saccharomyces cerevisiae*, we compared the UTRs captured by dRNA to those previously obtained by Park et al. (Park et al. 2014), in which the authors used an enzyme to remove the 5’cap of the mRNAs before ligating a specific adapter, and a threshold of 8 adenines before the 3’adapter to define the polyA tail. We could identify 99.1% of the 5’UTR and 98.9% of the 3’UTRs found in the previous study (Figure 1e), and recovered 1,185 additional 5’UTR and 916 3’UTRs. The size distribution of the 5’UTRs and 3’UTRs from both studies was similar (Figure S1), validating the approach.

The translation of uORFs is sometimes initiated at ATG near-cognate codons, or NTGs, namely TTG, CTG and GTG (Chen et al. 2020). Inspection of the frequency of these triplets in the 5’UTR and 3’UTR sequences revealed that the canonical translation start codon, ATG, was strongly underrepresented in 5’UTRs when compared to 3’UTRs (Figure 1f). In contrast, alternative NTG codons showed a similar low abundance in 5’UTR and 3’UTRs. This would be consistent with an effect of selection in limiting the number of ATG codons, but not the other NTG codons, in the 5’UTR. Translation starting at these codons is less efficient than translation from the ATG (Clements et al. 1988), and thus less selective pressure to remove them could be expected.

### Around one third of *S. cerevisiae* mRNAs contains translated ncORFs

The large number of available Ribo-Seq experiments performed in *S. cerevisiae* allows for a deep characterization of non-canonical translation in this species. We took advantage of an already compiled collection of high-quality Ribo-Seq datasets (Papadopoulos et al. 2024) to identify translated ORFs in mRNAs. After quality control, we used 640 million mapped Ribo-Seq reads from 100 different Ribo-Seq experiments. We predicted translated ORFs with RibORF v.2.0 (Ji et al. 2015)(Figure 2a). This software predicts ORFs with significant translation signatures on the basis of three-nucleotide periodicity and homogeneity of the Ribo-Seq reads along the ORF. Based on the results of the program, we grouped the ORFs in different groups: ‘not translated”, where there were 10 or fewer reads mapping to the ORF, ‘background’, when there were more than 10 reads but no significant translation signatures and, ‘translated’, with more than 10 reads and significant translation signatures (RibORF score cut-off of 0.6).

**Figure 2.**
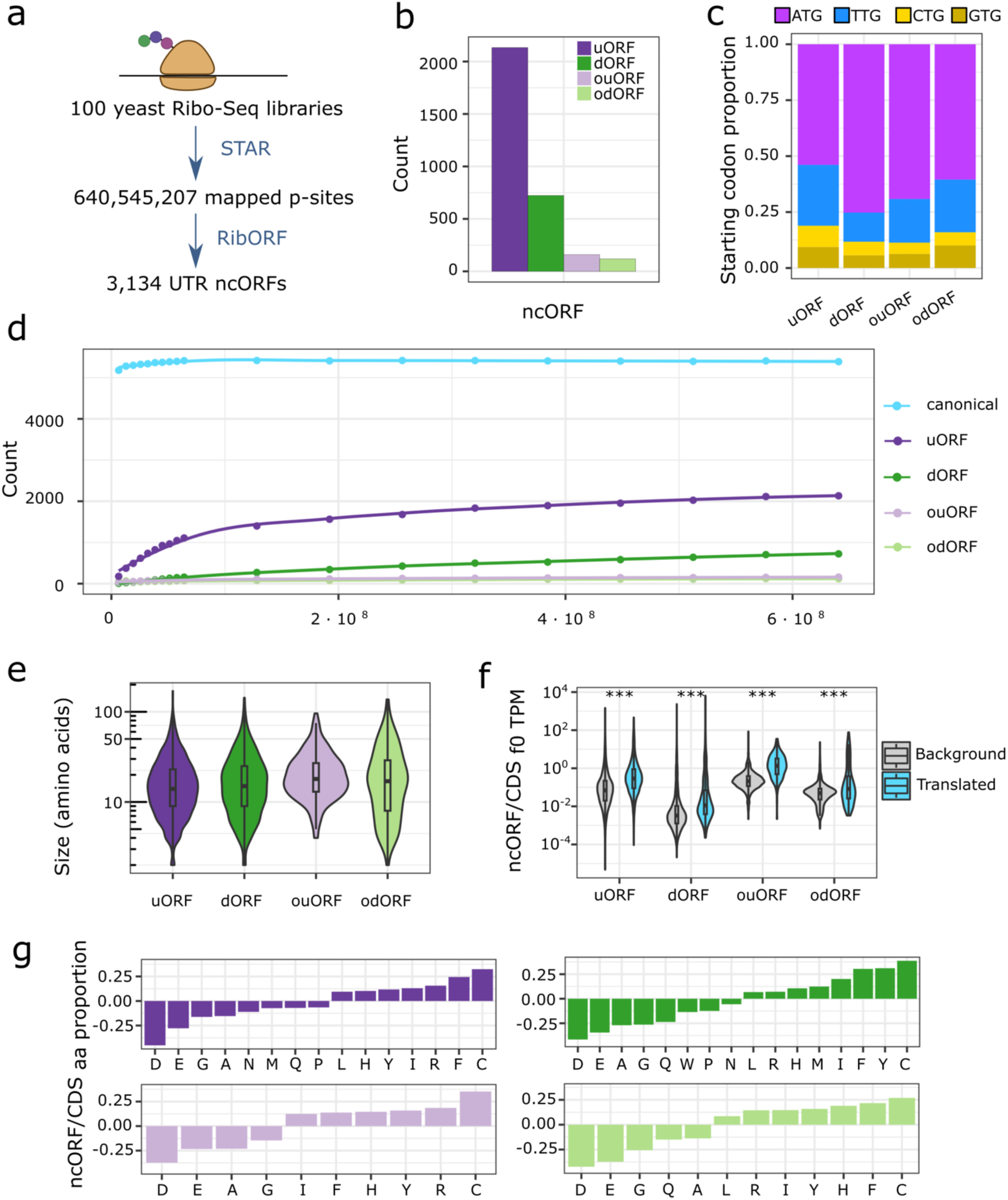
Main features of ncORFs in *S. cerevisiae* mRNAs. **(a)** Scheme of the procedure to analyze the ribosome profiling (Ribo-Seq) data. A 100 Ribo-Seq libraries were cleaned and mapped to the reference genome using STAR. Then we extracted the p-site per library depending on the length of the read with ribosomeProfilingQC. Finally, we used RibORF v. 2.0 to predict the translated ORFs (score ≥ 0.6). **(b)** Total number of ncORFs that were predicted to be translated. uORFs: 2,133; dORFs: 723; ouORFs: 159; odORFs: 119. **(c)** Frequencies of different start codons in translated ncORFs. **(d)** Number of detected translated ORFs for increasing numbers of subsampled sequencing reads. **(e)** Size in amino acids of the different types of ncORFs (see Table S6 for medians and quartiles). **(f)** Translation levels of different types of ncORFs relative to the CDS in the same transcript (see Table S7 for medians and quartiles). **(g)** Amino acid enrichment of ncORFs. Only cases associated with a false discovery rate under 0.01 are shown. Values over 0 indicate enrichment of that specific amino acid over the expectation for canonical coding sequences (CDS) in a logarithmic scale. Conversely, values under 0 indicate depletion of the indicated amino acid.

As expected, this analysis detected translation of the majority of annotated coding sequences (CDS), 5,389 out of the 6,685 (80.6%). Additionally, we detected significant translation of 2,133 uORFs, 723 dORFs, 159 ouORFs and 119 odORFs (Figure 2b, Tables S4 and S5). The proportion of uORFs that were translated *versus* all uORFs detected was 7.9%, whereas the same proportion for dORFs was 2.1% (Figure S2). The majority of translated ncORFs started at ATG, however, in the case of uORFs the TTG start codon was also observed in 27% of cases (Figure 2c).

The number of mRNAs which contained one or more translated uORF was 1,470 (24.2% of the genes with an annotated 5’UTR), whereas the number with at least one translated dORF was 625 (10.2 % of the genes with an annotated 3’UTR) (Figure S3). The values for ouORFs and odORFs were much lower, 159 genes (2.6%) and 119 genes (1.9%), respectively. In total, 2,179 mRNAs contained one or more translated ncORF.

We took advantage of the very high coverage of the Ribo-Seq data to inspect if the number of translated ORFs showed saturation with increasing number of sequencing reads, which would indicate that the catalogue was complete or nearly complete (Figure 2d, Figure S4). For canonical coding sequences, the saturation occurred very rapidly; we could recover 90% of translation events with only 1% of the reads. Saturation patterns were also observed for uORFs and ouORFs but were less clear for dORFs and odORFs. This would be consistent with the latter being translated at very low levels, and thus being less reproducible across experiments.

### uORFs are translated at higher levels than dORFs

The median size of the translated products of ncORFs ranged between 14 and 18 amino acids depending on the ORF type, compared to 412 amino acids for canonical proteins (Figure 2e, Table S6). For the ORFs classified as translated, the Ribo-Seq coverage is a direct measurement of the translation level, because each read corresponds to the ribosome-mRNA complex (Ingolia et al. 2009; Brar and Weissman 2015). We quantified ORF translation levels using exclusively in-frame reads (those for the correct frame, or f0); this allowed us to measure the translation of ouORFs and odORFs independently of the CDS. All ncORFs except ouORFs tended to show lower translation levels than the CDS in the same transcript (Figure 2f, Table S7). The translation levels were particularly low for dORFs and odORFs (Figure S5). Compared to canonical proteins, the microproteins translated from ncORFs tended to be enriched in cysteine, aromatic amino acids (F,Y,H) and arginine, and depleted of negatively charged amino acids (Figure 2g). These biases were similar for the different ncORF types.

### uORFs that are translated in different species are translated at levels comparable to the coding sequence

To investigate if the translation of the ncORFs in *S. cerevisiae* was phylogenetically conserved, we generated Ribo-Seq data for the other six *Saccharomyces* species shown in Figure 1a. The number of Ribo-Seq reads recovered per species ranged between 85 and 189 million reads (Table S8), and the reads showed very strong three-nucleotide periodicity (Figure S6), which is key to predict translated ncORFs. For *S. paradoxus* we combined these reads with previously published ones (Durand et al. 2019) to increase the depth. For predicting translated ncORFs in the UTRs, we used the gene annotations with 5’ and 3’UTRs obtained previously (see section “Reconstruction of full transcripts using Nanopore dRNA”). As for the *S. cerevisiae* data, we predicted translated ORFs in the six additional species using RibORF v.2.0 (Tables S9-S14). The number of predicted translated CDS ranged between 4,811 in *S. arboricola* and 5,321 in *S. kudriavzevii*. In addition, we predicted a total of 3,015 translated ncORFs across all six species, more than half of which were uORFs (1,653 cases)(Table S15).

To identify ncORFs with conserved translation across species, we first identified one-to-one orthologous genes between the seven species using proteinOrtho (Table S16). We then constructed pairwise mRNA alignments and identified orthologous ncORFs showing evidence of translation in *S. cerevisiae* and at least another *Saccharomyces* species. This resulted in the identification of 163 uORFs (7.64%) and 32 ouORFs (20.13%), but only 20 dORFs (2.77%) and 12 odORFs (10.08%)(Figure 3a, Tables S17 and S18). As expected, the species in which we observed more conserved translated events was *S. paradoxus*, which is the most closely related to *S. cerevisiae*. In addition, there were 125 ncORFs which showed translation in one or more of the three most distant species (*S. kudriavzevii*, *S. arboricola* and *S. uvarum*). Overall, there were 219 *S. cerevisiae* genes containing at least one ncORF translated in multiple species.

**Figure 3.**
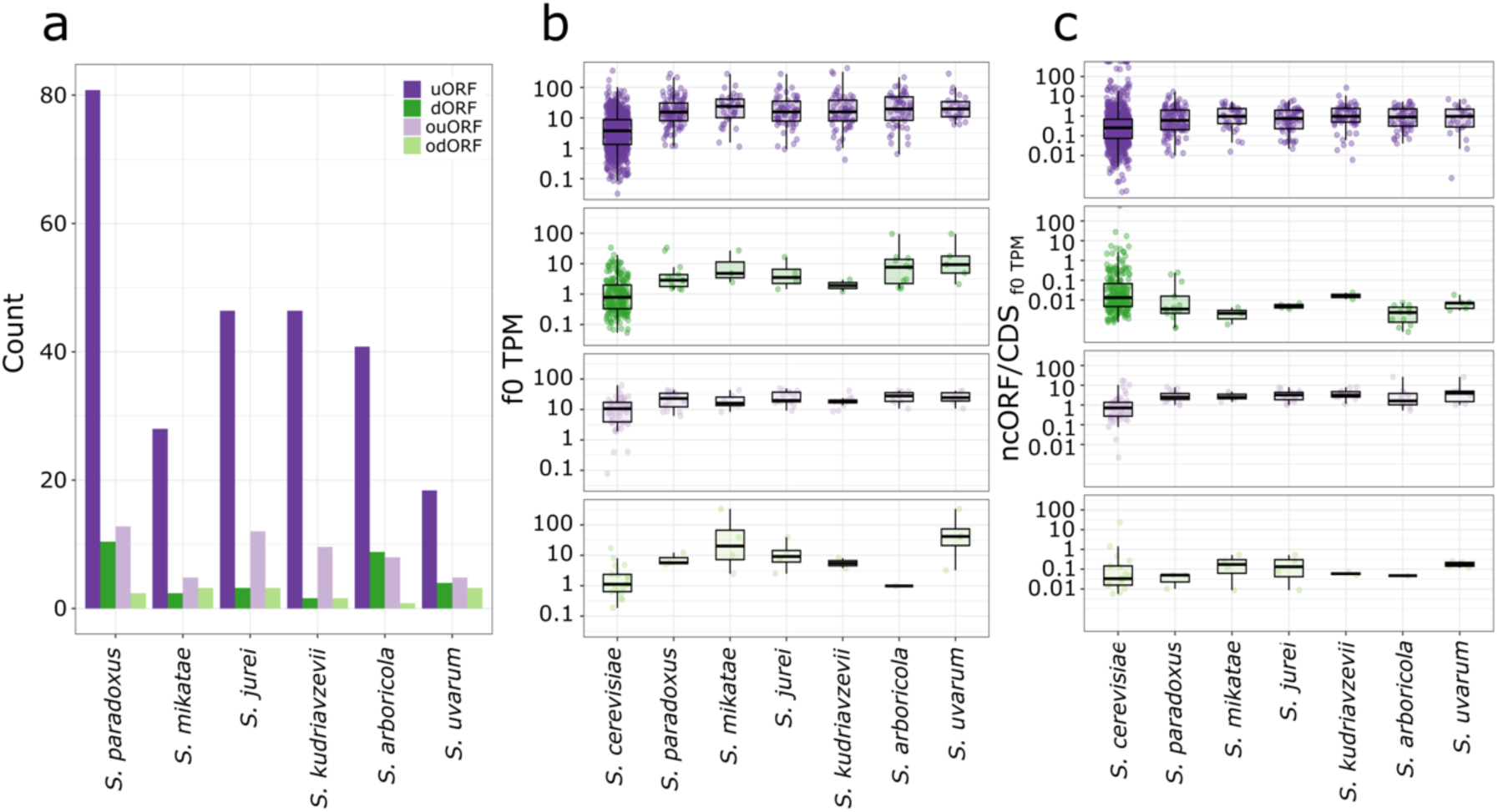
Features of conserved translated ncORFs in the *Saccharomyces* genus. **(a)** Number of ncORFs translated in *S. cerevisiae* and in other *Saccharomyces* species. **(b)** Absolute translation levels of ncORFs by conservation category (in TPM or transcripts per million), based exclusively on in-frame ribosome-profiling reads (f0)(see Table S19 for medians and quartiles). **(c)** Relative translation of ncORFs normalized to CDS expression, stratified by conservation category and considering only in-frame reads (see Table S20 for medians and quartiles).

In general, these ncORFs were translated at higher levels than the rest of *S. cerevisiae* ncORFs, especially in the case of uORFs and ouORFs (Figure 3b, Table S19). This was more clearly seen when compared to the CDS in the same transcript: the translation of conserved uORFs was, on average, similar to that of the CDS, and that of ouORFs slightly higher than that of the corresponding CDS (Figure 3c, Table S20). Instead, for dORFs and odORFs, the translation levels continued to be much lower than that of the CDS.

### Phylogenetically conserved translated uORFs show signatures of purifying selection at the protein level

The pool of ncORFs translated in several species provides a set in which to investigate possible functionalization events. We first tested if the conservation of some uORFs could be related to the control of translation of the main coding sequence in stress conditions. One example would be the uORFs located in the 5’UTR of the stress master regulator *GCN4* gene, which inhibit the translation of the main protein product in non-stress conditions (Hinnebusch 2005). In stress conditions, changes in the translation of the uORFs result in an increase in the translation efficiency (TE) of the *GCN4* canonical ORF. In a previous study (Moro et al. 2021), we systematically identified genes which showed a significant increase in TE during stress using previously published RNA-Seq and Ribo-Seq data from normal and oxidative stress conditions (Gerashchenko et al. 2012). This list included the previously described genes with regulatory uORFs *GCN4* and *HAP4*, which were translated in several Saccharomyces species. However, in general, the proportion of genes with increased TE in stress was similar among those containing *S. cerevisiae*-specific translated uORFs and those with phylogenetically conserved translated uORFs (14.4% *versus* 14.6%, respectively, Table S21). As the conservation of these uORFs is not correlated with translation regulation of the downstream CDS, an alternative explanation is required. One possibility is that many of them encode microproteins that have their own inherent functions. We investigated the functions of the genes containing them, which refer to functions of the main protein product. We found that the Gene Ontology function “transmembrane transporter activity” was strongly overrepresented (25 out of 157 genes, adjusted p-value = 2-13e-05)(Figure S7). They hydrophobic nature of the uORF-encoded microproteins suggests that some of them might also be located in the membrane.

The analysis of the rate of non-synonymous to synonymous substitutions (dN/dS) in aligned coding sequences is a good proxy for estimating purifying selection. Because of their small size, individual uORFs contain too few nucleotide substitutions for an accurate prediction of dN/dS, but several pairwise alignments can be concatenated and analyzed together. Values around 1 would indicate that the sequences are evolving neutrally, whereas values lower than 1 can be interpreted as a signature of purifying selection. In the case of annotated coding sequences typical dN/dS values are in the range 0.05-0.2 (Gonçalves et al. 2011), in accordance with strong purifying selection. For uORFs, we randomly selected 1/3 of the individual pairwise alignments and calculated the dN/dS of the concatenated alignment. We repeated this procedure 100 times, obtaining a distribution of dN/dS values. The median dN/dS values varied depending on the phylogenetic distance; they were around 0.8 for comparisons against *S. paradoxus* and *S. mikatae*, around 0.6 for *S. jurei*, *S. kudriavzevii* and *S. arboricola*, and 0.47 for the most distant comparison, *S. cerevisiae*-*S.uvarum* (Figure 4a, Table S22).

**Figure 4.**
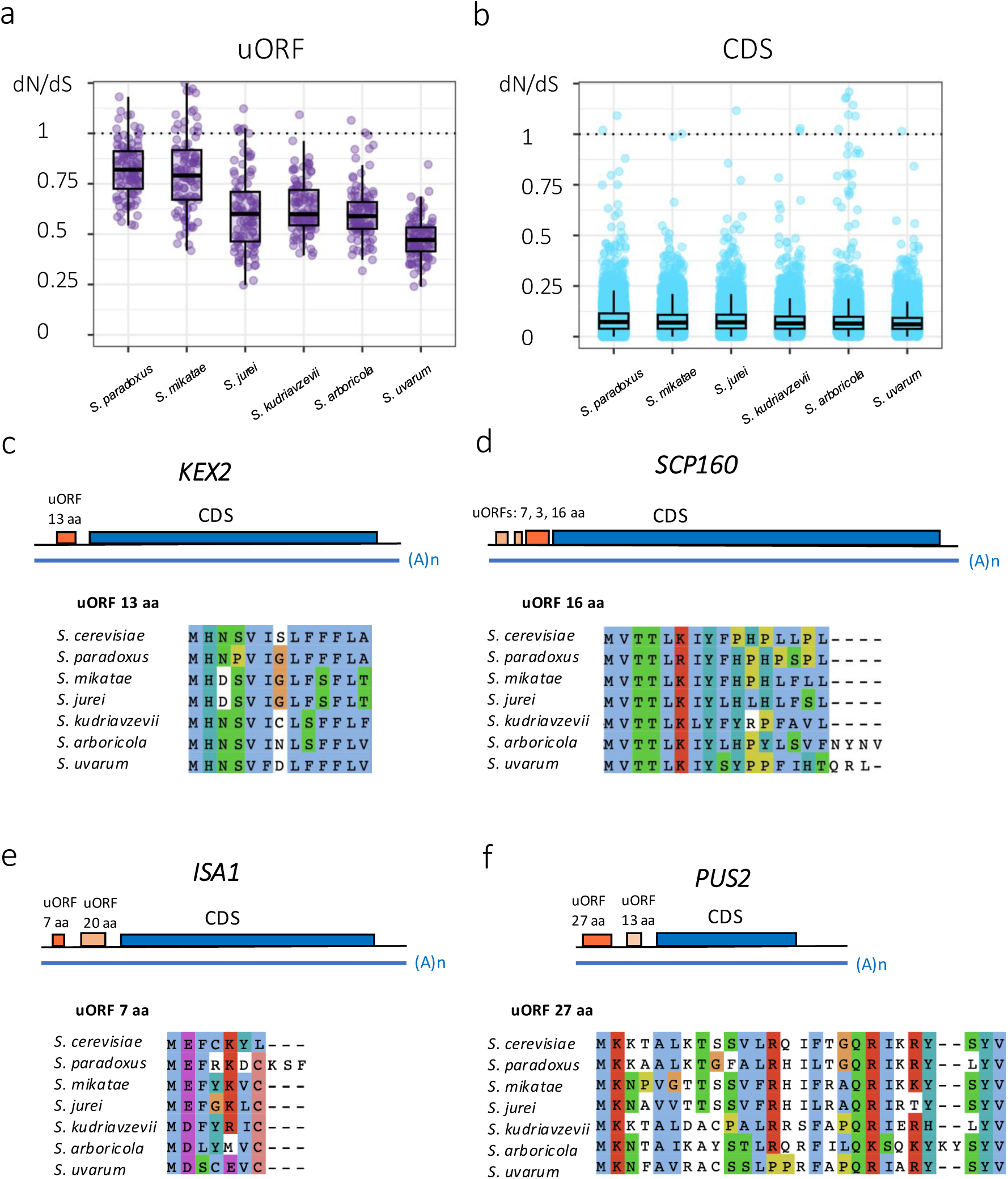
Constrained evolution of microproteins encoded by translated uORFs in the *Saccharomyces*. (a) Non-synonymous to synonymous substitution rate ratio (dN/dS) for uORFs translated in another *Saccharomyces* species. We randomly selected approximately 1/3 of the uORFs and concatenated the alignments to calculate dN/dS, repeating the procedure 100 times. dN/dS was estimated with the KaKs calculator (NG mode). The plot shows the distribution of the dN/dS values of the 100 concatenates (see Table S22 for medians and quartiles). **(b)** dN/dS distribution for CDS pairwise alignments (see Table S23 for medians and quartiles) for the same set of species. **(c)** Microprotein translated from a uORF in *KEX2* (YNL238W). The uORF encoding a microprotein of 13 aa in *S. cerevisiae* was significantly translated in *S. kudriavzevii*, *S. arboricola* and *S. uvarum*. **(d)** Microprotein translated from a uORF in *SCP160* (YJL080C). The uORF encoding a 16 aa protein in *S. cerevisiae* was significantly translated in *S. paradoxus*, *S. jurei* and *S. uvarum*. Two other uORFs located more upstream (7 and 3 aa) were only significantly translated in *S. cerevisiae*. **(e)** Microprotein translated from a uORF in *ISA1* (YLL027W). The uORF encoding a 7 amino acid (aa) protein in *S. cerevisiae* was significantly translated in all species except *S. uvarum*. A second uORF located more downstream (20 aa microprotein) was only found translated in *S. cerevisiae*. **(f)** Microprotein translated from a uORF in *PUS2* (YGL063W). The uORF encoding a microprotein of 27 aa in *S. cerevisiae* was significantly translated in *S. kudriavzevii* and *S. arboricola*, and the sequence conserved also in other species. A second downstream uORF encoding a 13 aa microprotein was only found translated in *S. cerevisiae*.

The observed increase of constraints in the microproteins sequences with phylogenetic distance might be expected. In the case of the *S. cerevisiae* - *S. paradoxus* pair, and unlike in other comparisons, the majority of the the uORFs were uniquely found in *S. paradoxus* (41 out of 78, 52.6%, Table S17), and therefore many of them were young. It is possible that the functions of some of these microproteins are not yet well-established and/or that, due to their very recent origin, the sequences are more free to evolve. For comparison, we calculated dN/dS at the CDS for the same pairs of species. In this case we obtained median dN/dS values in the range 0.06-0.07 for the different comparisons (Figure 4b, Table S23). The higher homogeneity of dN/dS values in this set was consistent with the fact that the majority of the *S. cerevisiae* protein coding genes (77.8%, 3,985 out of 5,124) had 1:1 orthologous across all the species compared so, for the most part, we were comparing the same set of genes. Taken together, we detected selection at the level of the microproteins encoded by uORFs that are translated in several species, however, the selective constraints are generally more relaxed than for standard proteins.

By examining the uORF sequence alignments, we identified cases with very strong amino acid conservation throughout the sequence, as well as cases in which there was substantial amino acid sequence variability across species. One example of the first class was a sequence encoding a 13 amino acid (aa) peptide in the gene *KEX2* (Figure 4c). This gene encodes a calcium-dependent serine protease involved in the activation of proproteins of the secretory pathway (Fuller et al. 1989). The uORF-encoded peptide was highly hydropohobic and well-conserved across all *Saccharomyces* species, with significant translation signatures identified in *S. kudriavzevii*, *S. arboricola* and *S. uvarum*. This consistent translation signal, taken together with the conservation of the sequence in all the analyzed species, strongly suggests that the uORF encodes a functional microprotein. Other examples of genes containing highly conserved translated uORFs were *SCP160*, an RNA-binding G protein effector of mating response (Frey et al. 2001; Guo et al. 2003), *ISA1*, encoding a protein required for the maturation of iron-sulfur cluster in the mitochondria (Mühlenhoff et al. 2011), or *MDL1*, encoding a mitochondrial membrane transporter (Young et al. 2001)(Figures 4d, 4e and S8).

The gene *PUS2*, encoding a mitochondrial tRNA:pseudouridine synthase (Behm-Ansmant et al. 2007), contained two translated uORFs encoding microproteins of length 27 and 13 aa, the longest of which was also found translated in other *Saccharomyces* species (Figure 4f). The sequence of this microprotein was quite variable across the species, although several positions were completely conserved. We also identified species-specific C-terminus extensions generated by stop-loss mutations in *S. kudriavzevii* (11 extra aa) and *S. arboricola* (16 extra aa)(Figure S9). These example illustrate that fact that microproteins can accommodate a substantial amount of sequence variability.

### The translation of uORFs is associated with the generation of shorter transcript isoforms

The 5’UTR of the *PHO80* gene contained two relatively long uORFs, encoding microproteins of sizes 18 and 30 amino acids in *S. cerevisiae* (Figure 5a). The translation of the uORFs was conserved as far as *S. kudriavzevii*, but missing from the two more distant species, *S. arboricola* and *S. uvarum*, suggesting that the translation began in the common branch of *S. cerevisiae* and *S. kudriavzevii*. *PHO80* encodes a cyclin that is involved in the response to nutrient levels and other limiting environmental conditions (O’Neill et al. 1996). The strong sequence conservation of the uORF-encoded microproteins observed in the multiple sequence alignments is consistent with conserved protein function.

**Figure 5.**
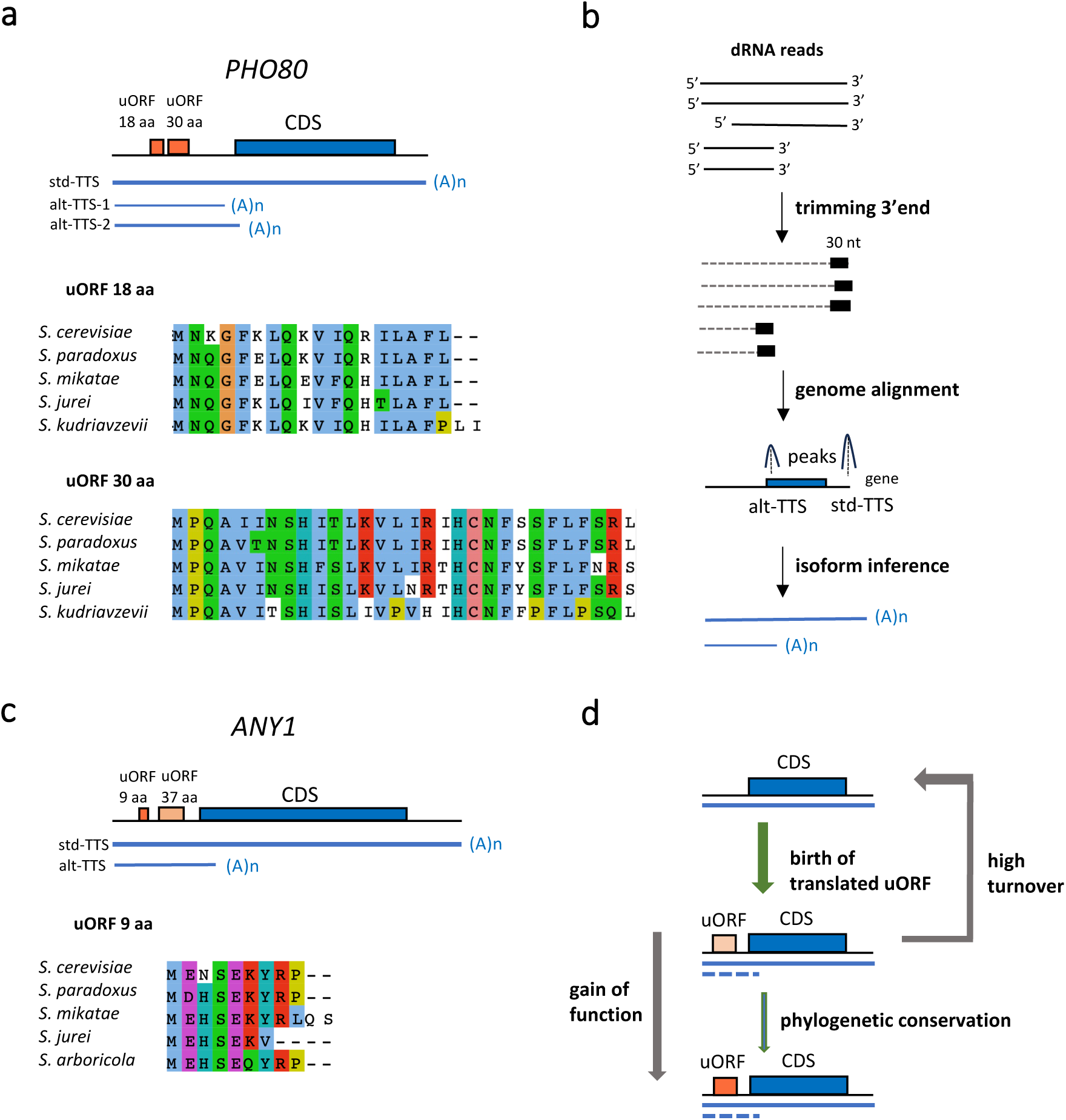
Phylogenetically conserved translated uORFs, alternative isoforms and evolutionary model. **(a)** Two translated uORFs in the *S. cerevisiae PHO80* gene (YOL001W), encoding proteins of 18 and 30 aa, were highly conserved in different Saccharomyces species. The first one was found to be translated in *S. paradoxus*, *S. mikatae*, *S. jurei* and *S. kudriavzevii*, and the second one in the same species except *S. mikatae*. Additionally, alignment of the dRNA reads to the genome revealed the presence of several transcript isoforms in the *S. cerevisiae* gene caused by alternative transcription termination sites (alt-TTS). **(b)** Computational pipeline to systematically detect transcription termination sites (TTS) using the dRNA reads. After genome alignment and read trimming, peaks were identified and isoforms reconstructed, using the positions of the TTS as the 3’end of the isoform. The more downstream TTS identified was considered to be the standard TTS (std-TTS). Alternative TTS (alt-TTS) corresponded to additional peaks that fell in the 5’UTR or the first 10% of the CDS, and which corresponded to isoforms of at least 50 nucleotides. Only TTS supported by at least 50 reads were kept. By definition the isoforms had different 3’end but the same 5’ end. The median size of the poly(A) tail for each isoform, denoted as A(n) in the Figure, was inferred from the information of the individual dRNA reads. **(c)** The gene *ANY1* (YMR010W) from *S. cerevisiae* contained two translated uORFS, encoding proteins of 9 and 37 aa. The first one was translated in *S. arboricola*, and the sequence also conserved in *S. paradoxus*, *S. mikatae*, *S. jurei* and *S. kudriavzevii*. We found evidence of an alternative isoform containing the two uORFs but not the CDS. **(d)** Evolutionary model of uORFs. High levels of uORF translation were observed using high coverage Ribo-Seq data for *S. cerevisiae*. Most uORFs were translated at low levels when compared to the CDS. Genes containing translated uORFs were enriched in alt-TTS generating isoforms that did not include the CDS. A subset of the translated uORFs was also found to be conserved and translated in other *Saccharomyces* species. These conserved uORFs were translated at much higher levels and displayed signatures of purifying selection at the protein level, indicating that they have acquired functions.

Intriguingly, visual inspection of the dRNA read mapped to the *PHO80* locus revealed a cluster of reads in the 5’UTR region, in addition to the usual reads ending at the 3’end of the mRNA. This suggested the presence of alternative shorter transcript isoforms. To identify such alternative isoforms in a systematic way in the complete set of genes, we extracted the 30 nucleotides located in the 3’end of all *S. cerevisiae* dRNA reads and performed peak calling with the program MACS2 (Zhang et al. 2008). This allowed us to approximately define transcription termination sites (TTS) using only the dRNA information (Figure 5b). We then selected TTS located in the 5’UTR or the 5’-most 10% of the CDS that resulted in isoforms with a minimum length of 50 nucleotides and supported by at least 50 reads (alt-TTS). For the *PHO80* gene, we obtained two alt-TTS, resulting in transcript isoforms of 488 and 561 nucleotides (alt-TTS-1 and alt-TTS-2). Both these isoforms contained the two conserved uORFs but not the CDS. The number of dRNA reads associated with alt-TTS-1 was 56, and for alt-TTS-2 108, compared to 895 reads for the longest isoform containing the CDS (std-TTS). This means that, together, the alternative isoforms represent 15.5% of the transcripts of the gene. Recovery of the polyA tail of the three isoforms from the dRNA reads confirmed that these were *bona fide* transcript 3’ends, with similar length poly-A tails (49.86 for alt-TTS-1, 42.81 for alt-TTS-2 and 45.95 for std-TTS).

In the complete gene set we identified 204 genes that had TTS peaks meeting our aforementioned criteria (Tables S24 and S25). In this set, the proportion of genes with translated uORFs was disproportionately high (95 out of 204, or 46.5%), nearly double that the random expectation (24.4% of genes containing translated uORFs). However, there were no significant differences between genes with *S. cerevisiae*-specific uORFs (81 out of 1,313, 6.17%) and genes with phylogenetically conserved uORFs (14 out of 157, 8.9%). One example in the latter class was *ANY1*, involved in endosome biogenesis (Gao et al. 2025). This gene contained three translated uORFs in S. cerevisiae, one of which was conserved down to *S. arboricola* (Figure 5c). An alt-TTS located at the beginning of the CDS resulted in a shorter 461 nucleotide transcript, which was represented by 643 dRNA reads, compared to 1,873 for the standard isoform. Thus, about one quarter of the dRNA reads corresponded to the alternative isoform containing only the uORFs. Another example was the *SER3* gene, which encodes a 3-phosphoglycerate dehydrogenase and alpha-ketoglutarate reductase (Albers et al. 2003). There were 4 translated uORFs in *S. cerevisiae SER3*, the first one of which was also significantly translated in *S. jurei* (Figure S10). The alternative isoform, containing all four uORFs, was 536 nucleotides long and it encompassed 2,256 reads, more than the 1,878 reads assigned to the standard isoform.

Taken together, the data shows that translation of uORFs in yeast mRNAs is very common, affecting approximately 1 out of 5 genes. Many of the uORFs are translated at very low levels, and this activity might not be sufficiently deleterious to be eliminated by natural selection (Lynch and Marinov 2015). It can be expected that the set of uORFs that is being translated change very quickly over time (Figure 5d). A subset of the uORFs, however, is translated at levels that are comparable to those of the CDS, and the translation signal can be detected in several *Saccharomyces* species. As indicated by the reduced number of non-synonymous substitutions compared to what would be expected by chance, the microproteins encoded by these uORFs are very likely to be functional. They are encoded by polycistronic transcripts and in some cases they might interact with the main protein product. Intriguingly, we found evidence that some of them can also be translated by transcripts that do not include the CDS. This could be a way to further regulate the relative abundances of the different protein products of the gene.

## DISCUSSION

Genome-wide scans of translation using Ribo-Seq have shown that the translation of ncORFs in the UTRs of eukaryotic genes is widespread (Ingolia et al. 2009, 2011; Duncan and Mata 2014; Mudge et al. 2022; Chothani et al. 2022). However, without comparable Ribo-Seq data from several closely related species, it is difficult to evaluate if this activity is of any functional significance. To fill this gap, we took advantage of the *Saccharomyces* fungi as a model eukaryotic genus to study the translation of uORFs and dORFs across evolutionary time. Our approach combined the exhaustive identification of translated ncORFs in *S. cerevisiae* with the generation of new dRNA and Ribo-Seq data for six other species. We uncovered tens of microproteins encoded by uORFs that were translated in several species and which showed hallmarks of functionality, such as high translation levels and significant sequence conservation across species.

The reconstruction of the gene 5’ and 3’UTR sequences was a prerequisite for the subsequent characterization of the ncORF translatome. Whereas information on 5’ and 3’ ends of the transcripts can be retrieved from the literature in the case of *S. cerevisiae* (Moqtaderi et al. 2013; Park et al. 2014; Zhang and Dietrich 2005b), this data was completely lacking for other species of the genus. The high coverage of the dRNA reads allowed us to reconstruct UTRs for the vast majority of the annotated genes in each species. In the case of *S. cerevisiae*, the number of reconstructed 5’ and 3’UTRs surpasses those in the study of Park et al. (2014). The gene annotations generated here will be useful for future comparative studies. In addition, similar approaches could be used to reconstruct more complete transcriptomes of fungi species outside the *Saccharomyces* group.

We observed an important asymmetry in translation activity between the 5’UTR and the 3’UTR, with the former containing a much larger number of translated ncORFs than the latter. The largest class of such ORFs (uORFs), represented about two thirds of all translated ncORFs in *S. cerevisiae* (2,133 out of 3,134). This extensive translation of uORFs happens despite evidence of negative selection for the formation of the ATG triplet. In humans, the analysis of single nucleotide polymorphism data has provided evidence that the formation of new uORFs can be deleterious and cause disease (Whiffin et al. 2020). A survey of different species found that the observed *versus* expected ratio of uORFs initiating at ATG was generally lower than 1, denoting avoidance of the ATG triplet (Zhang et al. 2021). In agreement with these earlier findings, here we found that the ATG triplet was rarely found in the 5’UTR when compared to the 3’UTR. Because the 5’UTR is more accessible to the scanning ribosome than the 3’UTR, this would be consistent with selection against the translation of uORFs.

The majority of the ncORFs detected in *S. cerevisiae* were translated at much lower levels than the CDS. This probably means that, in many cases, the activity is residual, and that the ribosomes normally bypass the uORFs. Such low translation levels might not be sufficiently deleterious to be eliminated by selection (Lynch and Marinov 2015). The generation of Ribo-Seq data from the other six species was key to uncover a group of ncORFs that was translated at much higher levels and which showed conserved translation across several species. These microproteins are very likely to have acquired functions. Taken together, the data supports a model in which pervasive ncORF provides a reservoir of microproteins that can be tested for new functions. A subset of these translation events will be preserved by selection. This is similar to current models for *de novo* gene birth, in which small ORFs are initially translated by accident from ‘non-coding’ transcripts, with a few surviving and being maintained by selection (Toll-Riera et al. 2009; Carvunis et al. 2012; Reinhardt et al. 2013; Ruiz-Orera et al. 2018; Schmitz et al. 2018; Van Oss and Carvunis 2019; Blevins et al. 2021; Durand et al. 2019). We found that the microproteins encoded by ncORFs tend to be highly hydrophobic and/or positively charged. The same tendencies are observed for *de novo* proteins, and they can be explained by an origin from previously non-coding sequences (Montañés et al. 2023).

Previous studies used uORF sequence conservation across species to detect putative functional candidate sequences (Cvijović et al. 2007; Selpi et al. 2009; Zhang et al. 2018, 2021; Zhang and Dietrich 2005a). The effects on the translation of the downstream CDS were in some cases tested using luciferase reporter systems. For example, using this methodology, Zhang and Dietrich (2005) identified five cases in which the uORFs had a regulatory effect on the CDS. For two of these cases, *FOL1* and *MKK1,* we identified conserved translated uORFs in other species, although these genes were not in the list of genes showing increased TE in oxidative stress using previously published data (Gerashchenko et al. 2012; Moro et al. 2021). We also found conserved translation for two classical examples of regulatory uORFs in yeast, *GCN4* and *HAP4* (Grant et al. 1995; Hinnebusch 2005; Vilela and McCarthy 2003). In this case, the two genes showed higher TE in oxidative stress than in normal conditions.

Overall, we identified 163 uORFs in *S. cerevisiae* whose translation was also detectable in other species. More than half of them (85) showed deep translation conservation in *S. kudriavzevii*, *S. arboricola* or *S. uvarum*. We propose that many of these microproteins are likely to be functional *per se*, beyond possible effects of the uORF translation activity on the CDS. Consistent with this, we observed that the number of non-synonymous substitutions in these uORFs was consistently lower than expected. Inspection of multiple sequence alignments provided further evidence of the conservation of the amino acid sequences. Some microproteins, such as a 13 aa peptide encoded by a uORF in *KEX2* and a 16 aa peptide in *SCP160*, were highly conserved throughout their sequence, whereas others, including the 27 aa peptide in *PUS2*, showed an overall more variable sequence, although with some very well conserved positions.

Several examples of polycistronic genes encoding microproteins translated from the 5’UTR have been described in mammals (Samandi et al. 2017; Ajala and Vanderperre 2025). For example, the mitochondrial elongation factor 1 gene (*MIEF1*) encodes a microprotein translated from an uORF, which has been shown to regulate mitochondrial translation (Rathore et al. 2018). According to a proteogenomics study, the MIEF1 70 aa microprotein is more abundant than the canonical 463 amino acid MiD51 protein (Delcourt et al. 2018). Another example is a uORF-encoded microprotein in the gene *ASNSD1*, which is required for medulloblastoma cell survival (Hofman et al. 2024). The micropeptide STMP1, encoded by an uORF, is associated with metastasis and recurrence of hepatocellular carcinoma (Xie et al. 2022). A microprotein of 31 aa, MP31, is translated from a uORF in the *PTEN* gene. This protein is involved in regulating mitochondrial lactate oxidation (Huang et al. 2021).

The functions of uORF-encoded microproteins might be related to the main protein product or be unrelated. Analysis of co-immunoprecipitated proteins for several candidates in humans identified 5 cases which formed complexes with the main protein product, including the *MIEF1* microprotein, along with two others that appeared to have independent functions (*TBPL1, ARL5A*)(Chen et al. 2020). To our knowledge, in yeast there are no examples of functional microproteins encoded by uORFs. Well-studied cases of uORFs whose translation has a regulatory role on the main protein product are typically very short. For examples, the four uORFs is *S. cerevisiae GCN4* are only 2-3 codons long (Mueller and Hinnebusch 1986). We found that some of the uORFs that show conserved translation across species and high amino acid conservation were relatively long, one example being the uORF-encoded 30 aa microprotein in *PHO80*. However, others were remarkably small, such as a 7 aa and 9 aa microproteins in *ISA1* and *ANY1*, respectively. Proteins in the range of 10-12 aa have been found to play cellular roles in insects and plants (Rohrig et al. 2002; Savard et al. 2006; Kondo et al. 2007; Zanet et al. 2015). Our study points to a much wider landscape of functional small proteins than previously suspected.

Several mechanisms have been proposed for the translation of multiple ORFs in the same transcript, including ribosome leaky scanning, translation reinitiation and internal ribosome entry sites (Andrews and Rothnagel 2014; Vilela and McCarthy 2003; Young and Wek 2016). In the first case, a subset of the ribosomes scanning the mRNA from the cap bypasses the uORF and start translation directly at the CDS. In the second case, the ribosome continues to scan the mRNA after translating the first ORF, being able to initiate translation when a new ORF/CDS is encountered. A special case is ouORFs, in which translation terminates after the start of the CDS. One possibility is that, after ending the translation of the ouORF, translating ribosomes are able to reinitiate translation of the CDS, provided the distance between the stop codon of the ouORF and the start codon of the CDS is of only a few nucleotides (Young, Baird, and Wek 2016). In this case the translation of the CDS would be expected to be less efficient than that of the ouORF. This is consistent with our observations of increased translation levels in the ouORF *versus* the CDS, especially in those cases conserved across species. Finally, although most yeast mRNAs follow a cap-dependent translation mechanism, the mRNAs encoding the transcription factors TFHD and HAP4 have been shown to be translated in a cap-independent manner (Iizuka et al. 1994).

One unexpected finding of the study was the identification of unannotated transcript isoforms that did not contain the CDS but only translated uORFs. An excess of alternative polyadenylation sites at the beginning of the CDS was noted in a previous study in yeast (Moqtaderi et al. 2013). In this study we extended the characterization of alternative TTS to the newly mapped 5’UTRs. The TTS positions were defined as “peaks” using the 3’ends of the dRNA reads aligned to the genome. After discarding TTS signals that could correspond to different, overlapping, genes, we identified cases falling in the 5’UTR or the first 10% of the CDS (108 and 106 cases, respectively). Nearly half these cases contained translated uORFs. The relative abundance of the different isoforms could be quantified using the dRNA reads. We found cases in which the alternative transcript was more abundant than the standard one, such as *SER3*. Alternative isoforms linked to the expression of uORFs have also been described in the *McKusick–Kaufman syndrome* (*MKKS*) gene (Akimoto et al. 2013). This mechanism may have evolved to fine tune the levels of translation of the different proteins encoded in the same gene.

In conclusion, this study significantly expands our view on the breadth and evolutionary conservation of ncORF translons, uncovering tens of novel microproteins that are encoded by polycistronic transcripts and which show deep conservation in the *Saccharomyces*. This novel catalog is expected to enhance future investigations into the functions of microproteins and their expression regulation.

## METHODS

### Yeast cultures

*Saccharomyces cerevisiae* S288C, *Saccharomyces paradoxus* NRRL Y17217, *Saccharomyces mikatae* IFO1815, *Saccharomyces jurei* D 5088 NCYC 3947, *Saccharomyces kudriavzevii* IFO 1802, *Saccharomyces arboricola* H-6 and *S. uvarum* CBS7001 were grown in in YPD medium at 30 °C to mid-log phase (OD₆₀₀ = 0.5–0.8).

### Direct RNA sequencing (dRNA)

In order to obtain RNA, yeast cultures (25–50 mL) were centrifuged at 1500 rpm for 3 min and washed with H2O, and cell pellets were immediately kept on ice. Each sample was then resuspended in 0.4 mL of AE buffer (50 mM sodium acetate at pH 5.3, 10 mM EDTA at pH 8.0). Sodium dodecyl sulfate was then added to a final concentration of 1%, and proteins and DNA were extracted by adding 0.6 mL of acidic phenol/chloroform (V/V), followed by incubation for 5 min at 65°C. The aqueous phase was separated by centrifugation at 14,000 rpm for 2 min at 4°C, washed with chloroform and separated by centrifugation at 14,000 rpm for 2 min at 4°C. RNA was precipitated from the aqueous phase with ethanol. We subsequently performed poly(A)+ RNA purification using the NEBNext Poly(A) magnetic isolation module and concentration with the Monarch RNA cleanup kit. The poly(A)+ purification steps were performed at the Genomics Core Facility of Universitat Pompeu Fabra.

The poly(A)+ RNA was used for dRNA-seq in an ONT Gridion X4. dRNA-seq offers the advantage over cDNA sequencing in that strand orientation information is maintained. The protocol involves adaptor ligation, and the molecules pass through an ionic current, adaptors and poly(A)+ tail first and then the rest of the molecule. The samples from each species were run in four flowcells. For each run, we used ∼600 ng of poly(A)+ RNA in 10 μL of volume. The dRNA-seq kit SQK-RNA002 was used. The base-calling was performed on live mode through Dorado v7.3.11 integrated on MinKNOW v24.02.16. Nanopore dRNA-seq and base-calling was performed by the Centro Nacional de Análisis Genómico (CNAG). We pulled together the output of four runs, obtaining a total of 33-49 dRNA million reads per species (Table S1). We polished the dRNA reads using fmlrc (v1.0.0) (Wang et al. 2018), with Illumina short reads generated for the same species.

For each species, we mapped the dRNA reads to the reference genome using minimap2 (Li 2018). The genome sequences were extracted from Genbank at the National Center for Biotechnology Information (NCBI): saccharomyces_cerevisiae_R64-1-1.fsa (downloaded 12 April 2024), Saccharomyces_paradoxus_CBS432.fasta (downloade 12 April 2024), Saccharomyces_mikatae_IFO1815.fna (downloaded 12 April 2024), Saccharomyces_jurei_D5088NCYC3947.fasta (downloaded 30 Jan 2025), Saccharomyces_kudriavzevii_IFO1802.fna (downloaded 12 April 2024), Saccharomyces_arboricola_H6.fasta (downloaded 30 Jan 2025), Saccharomyces_uvarum_CBS7001.fna (downloaded 12 April 2024).

### Illumina RNA sequencing

We obtained total RNA from cultures of the seven species using the RNeasy kit (Qiagen) with enzymatic cell lysis, followed by the manufacturer’s instructions. Poly(A)+ purification was performed as for the RNA extracted to perform dRNA sequencing. Paired-end sequencing (2 × 150 bp) was performed on an Illumina platform at the Genomics Unit of the Centre for Regulatory Genomics (Barcelona, Spain). We obtained a total of 65-82 raw million reads per species (Table S1).

### Transcript reconstruction and annotation of 5’ and 3’UTRs

For the reconstruction of the 5’ and 3’ untranslated regions (UTRs), we first estimated the strand-specific coverage of the aligned direct RNA reads using BEDTools (v2.30). Coverage profiles, together with species-specific annotations, were used as input for an in-house Python script (https://github.com/JC-therea/Charles_tools/blob/main/utr.bam.extender.py) to identify transcription start sites (TSS) and transcription termination sites (TTS). The script detects coverage drops near annotated transcripts, applying a minimum threshold of 10% relative to the previously defined start or end and requiring at least 10 supporting reads. Newly identified UTRs were either trimmed or discarded if they partially or completely overlapped with the CDS of another gene. Finally, the annotation was harmonized using AGAT (https://www.doi.org/10.5281/zenodo.3552717). We reconstructed UTRs for over 5,000 genes in each of the seven species (Table S2). Gene annotations containing 5’UTR and 3’UTR coordinates for each of the seven species are available as supplementary material.

### Identification of translation signatures in *Saccharomyces cerevisiae* using 100 Ribo-Seq datasets

We used data from 100 *S. cerevisiae* Ribo-seq libraries (Table S26) to identify translation signatures in the ORFs. All the experiments corresponded to yeast grown in rich media. The P-site was individually estimated in each experiment using the bioconductor package ribosomeProfilingQC 1.23.0 (https://bioconductor.org/packages/ribosomeProfilingQC). Subsequently, all libraries were processed with ribORF v.2.0 (Ji 2018). Each library underwent individual trimming using cutadapt (Martin 2011). Subsequently, reads were aligned to the reference genome (NCBI assembly: GCA_000146045.2; annotation source: annotation-source SGD R64-3-1) employing STAR (Veeneman et al. 2016). We obtained a total of 640,545,207 mapped Ribo-Seq reads for experiments and read lengths showing at least 60% of periodicity. After trimming the reads by their offset with RibORF all the samples were merged in a single file with samtools. RibORF was also utilized to estimate the counts per ORF, configured with options −l 6 and -r 1 (Ji 2018). For estimating ORF translatability, we applied the recommended minimum cutoff (-r 11). We applied the default RibORF score cut-off of 0.6. We consider any ORF starting with NTG, as ncORFs are known to frequently start with non-ATG codons. Transcripts per Million (TPM) were calculated using the RibORF output and considering all the reads that mapped to canonical, uORF, ouORF, dORF and odORF regions. In cases where multiple ORFs overlapped, we selected the ORF with an ATG start codon; if none of the overlapping ORFs contained an ATG, we retained the longest ORF.

To investigate the relationship between the number of reads and ORF translation detectability, we subsampled the file containing the reads from all the samples with samtools (*samtools view -h -s*). For each datapoint, we performed the same operation 10 times and took the average value.

### Generation of Ribo-Seq data and identification of translated ORFs in six additional

#### Saccharomyces species

Flash frozen cell pellets were ground by mortar and pestle under liquid nitrogen in the presence of 2mL polysome lysis buffer (20mM Tris pH 7.5, 150mM KCl, 5mM MgCl2, 1mM DTT, 1% Triton X-100) supplemented with cycloheximide (100μg/mL). Lysates containing 30μg RNA, as measured using a Qubit 4.0 fluorometer, were diluted to 200μL in polysome lysis buffer. Lysates were digested in the presence of 15U RNase1 for 45 min at room temperature. RNA was subsequently purified from lysates before undergoing PNK end repair. Ribosome protected fragments were size selected on 15% urea PAGE gels. Enriched fragments were reverse transcribed into cDNA containing Illumina compatible adapter sequences before contaminating rRNA sequences were depleted using EIRNABio’s custom designed biotinylated depletion oligos for yeast samples. Samples were subsequently amplified by PCR for the generation of Illumina compatible cDNA libraries.

For stranded RNA-seq from the same set of samples, total RNA was extracted from 100μL of lysate using TRIzol, before being rRNA depleted using the QIAseq FastSelect rRNA Yeast Kit (#334215, Qiagen, Germany), fractionated, and converted into Illumina compatible cDNA libraries. Both RNA-seq libraries and Ribo-seq libraries were sequenced 150PE on Illumina’s NovaSeq X platform. The number of raw Ribo-Seq sequencing reads generated per species and used in this study is indicated in Table S8. In the case of *S. paradoxus* we also used already existing data from Durand et al. (2019) and from SRA entries SRR5996797, SRR5996801 and SRR948564. The total number of raw reads used for *S. paradoxus* was 667,705,764. Prediction of translated ORFs was performed with ribORF v.2.0 (score cut-off 0.6)(Ji 2018). The results of these predictions are available in Tables S9-S15.

#### Inter-species sequence comparisons

We used proteinOrtho to identify orthologous sequences across the species (*proteinortho - synteny*)(Lechner et al. 2011)(Tables S16 and S27). We then aligned one-to-one orthologous mRNAs, one from *S. cerevisiae* and the second from another *Saccharomyces* species, with the program needle from the EMBOSS package (Rice et al. 2000). We used the pairwise alignments to identify conserved ncORFs and CDS between *S. cerevisiae* and a second species. Conservation of ncORFs required that at least the start codon or the stop codon were in the same position in the alignment plus that the overlapping region covered represented 90% or more of the *S. cerevisiae* ORF. The data of conservation of ncORFs for pairs of species was integrated into a single table. We employed the Jalview program with Clustal coloring (Waterhouse et al. 2009) for the visualization of uORF-derived microprotein sequence alignments.

#### Measuring the relationship between phylogenetic conservation of uORFs and changes in the TE of the CDS during oxidative stress

We obtained a list of 816 genes whose translation is upregulated during oxidative stress (0.5 mM H2O2 for 30’) from Moro et al. (2021). These genes had been defined as having significantly higher translational efficiency (TE) in oxidative stress conditions than in normal growth conditions (816 out of 6692). TE was calculated using RNA-Seq and Ribo-Seq data from Geraschenko et al. (2012). The program RiboDiff (Zhong et al. 2017) was then used to identify genes that showed significantly increased TE in stress than in normal conditions (adjusted p-value 0.05). The proportion of genes with uORFs translated only in *S. cerevisiae* and which showed increased TE in stress was 167 out of 1,313 (12.7%). The proportion of genes with uORFs translated also in other species and which showed increased TE in stress was 23 out of 157 (14.6%). The latter class included the genes *GCN4* (YEL009C) and *HAP4* (YKL109W), known to contain regulatory uORFs.

#### Measuring purifying selection signatures

We calculated the rate of non-synonymous to synonymous substitutions (dN/dS) in the regions that corresponded to conserved ORFs in the pairwise alignments using the KaKs calculator v. 2.0 with NG mode (Wang et al. 2010). In the case of the CDS, we performed this calculation directly using the information from the mRNA alignments, trimmed to the CDS region. In the case of uORFs, we manually curated the alignments, eliminating pairwise alignments in which there was a frameshift mutation (indel of size not multiple of three) affecting most of the amino acid sequence (50 out of 262 alignments, representing 19.1% of the total). The final number of uORF alignments used for dN/dS calculation was the following: *S. cerevisiae* and *S. uvarum* 18, *S. cerevisiae* and *S. arboricola* 35, *S. cerevisiae* and *S. kudriavzevii* 35, *S. cerevisiae* and *S. jurei* 35, *S. cerevisiae* and *S. mikatae* 20, *S. cerevisiae* and *S. paradoxus* 69. Because the uORFs are too short to perform accurate dN/dS estimations, we randomly selected approximately 1/3 of the alignments and concatenated them before calculating dN/dS. The exact number of alignments in each case was the following: *S. paradoxus* 23, *S. mikatae* 6, *S. jurei* 11, *S. kudriavzevii* 11, *S. arboricola* 11, *S. uvarum* 6. We repeated this procedure 100 times, obtaining a distribution of dN/dS values for each comparison between *S. cerevisiae* and another species. In the case of dORFs, ouORFs and odORFs, the low number of cases in which the translatioin was conserved in other species, together with the short size of the ORFs, precluded the estimation of accurate dN/dS values.

#### Identification of alternative transcription termination sites (alt-TTS)

We trimmed the 30 nucleotides at the 3’end of the dRNA reads aligned to the genome with the python module pysam. Subsequently, we performed peak calling with the program MACS2 (Zhang et al. 2008). For each significant peak, we extracted the position with the highest coverage. This position represented a predicted TTS. If the gene had a single predicted TTS, we named it std-TTS. If the gene had several predicted TTS, the most downstream was the std-TTS and the other ones alt-TTS. Using this information, we calculated the length of the different isoforms, from the 5’end of the gene to the TTS. The poly(A) tail length of each isoform was the average poly(A) length of the different dRNA reads associated with the isoform. Subsequently, we selected alt-TTS that fell in the 5’UTR or the first 10% of the CDS, were 50 nucleotides long and corresponded to at least 50 reads, for further analysis.

## Data availability

Gene annotations for the seven Saccharomyces species including 5’UTR and 3’UTR genomic coordinates, as well as sequence pairwise alignments of uORFs with conserved translation are available from Figshare (https://doi.org/10.6084/m9.figshare.31400334). The sequencing data can be accessed at the Sequence Read Archive (SRA) BioProject entry PRJNA1429909. This includes Nanopore dRNA and mRNA Illumina sequencing data for *S. cerevisiae*, *S. paradoxus*, *S. mikatae*, *S. jurei*, *S. kudriavzevii*, *S. arboricola* and *S. uvarum, as well as the* Ribo-Seq sequencing reads for the last six species, together with RNA-Seq reads for the same samples.

## ACKNOWLEDGEMENTS

We are grateful to David Peris for strains Saccharomyces jurei D 5088 NCYC 3947 and Saccharomyces kudriavzevii IFO 1802 and Saccharomyces arboricola H6. Ribo-seq libraries for S. paradoxus, S. mikatae, S. jurei, S. kudriavzevii, S. arboricola and S. uvarum were generated and sequenced by EIRNABio (https://eirnabio.com). We acknowledge funding from Ministerio de Ciencia e Innovación Agencia Estatal de Investigación grant PGC2018-094091-B-I00 (cofunded by Fondo Europeo de Desarrollo Regional), as well as PID2021-122726NB-I00 funded by MCIN/AEI/10.13039/501100011033 and by “ERDF: A way of making Europe”, as well as PID2022-136939OBI00/MICIN/AEI/10.13039/501100011033 and “ERDF a way of making Europe”, co-funded by the “European Union”. We also acknowledge funding from Generalitat de Catalunya, grants 2021SGR00042 and 2021SGR00176. We also received funds from an institutional “Unidad de Excelencia María de Maeztu,” CEX2024-001431-M, MICIU/AEI/10.13039/501100011033. The work was also funded by the European Union (ERC, NovoGenePop, project number 101052538). Views and opinions expressed are however those of the authors only and do not necessarily reflect those of the European Union or the European Research Council. Neither the European Union nor the granting authority can be held responsible for them.

## AUTHOR CONTRIBUTIONS

JCM, CP and AS were involved in the preparation of samples for Illumina RNA sequencing. JCM, WRB and EH contributed to the isolation of RNA and the design of dRNA experiments. JCM, CP, AS, M T-P and JD were involved in the design and preparation of samples for the Ribo-Seq experiments. JCM annotated the 5’ and 3’UTR regions of mRNAs using dRNA data. JCM and CP performed bioinformatics analyses of the Ribo-Seq data. SA-O and JCM developed a pipeline for the identification of transcription termination sites using dRNA data. JCM, CP and MMA participated in the evolutionary analysis of the data, including the construction of sequence alignments and the analysis of natural selection signatures. MMA supervised the study and wrote the manuscript with input from all authors.

## Notes

### Competing Interest Statement

The authors have declared no competing interest.

