## Supplementary Figures for "Evolutionary emergence and preservation of microproteins encoded by upstream ORFs"

Montañés JC, Papadopoulos C et al.

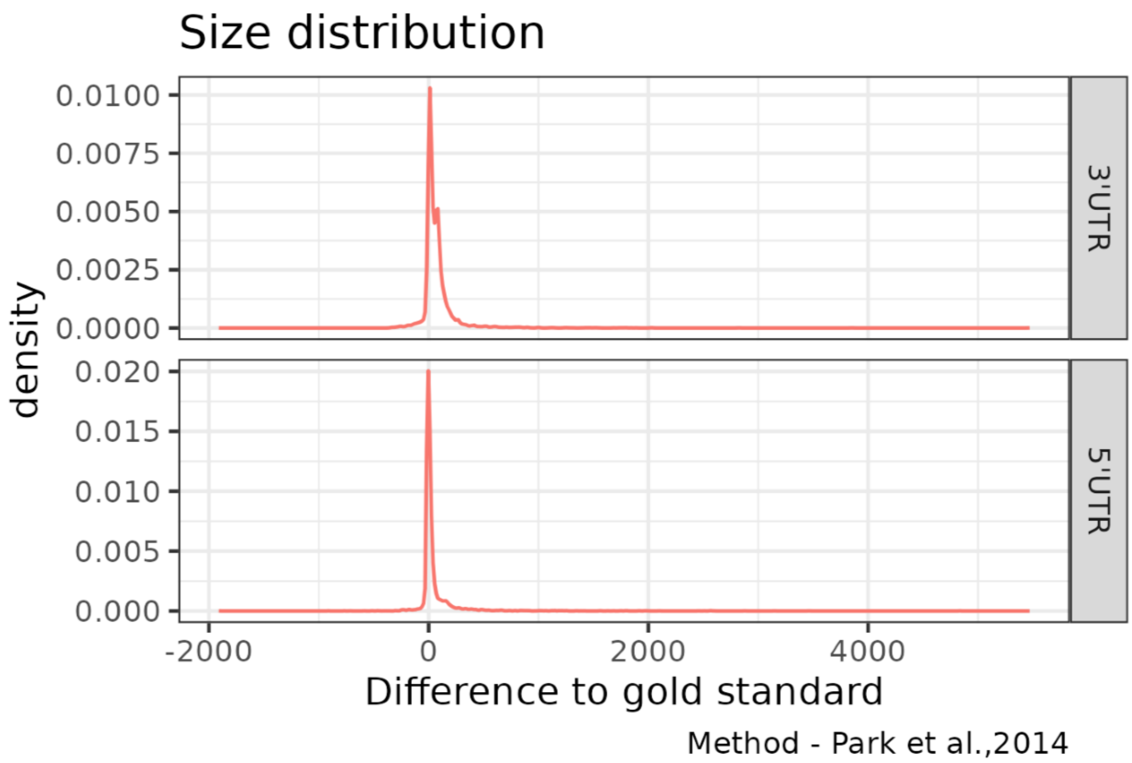


**Figure S1. Comparison of UTR sizes between our dRNA-based pipeline in *S. cerevisiae* and Park et al. 2014.**The distribution of the difference in size (nucleotides) between our method and the estimation by Park et al., 2014 is shown.

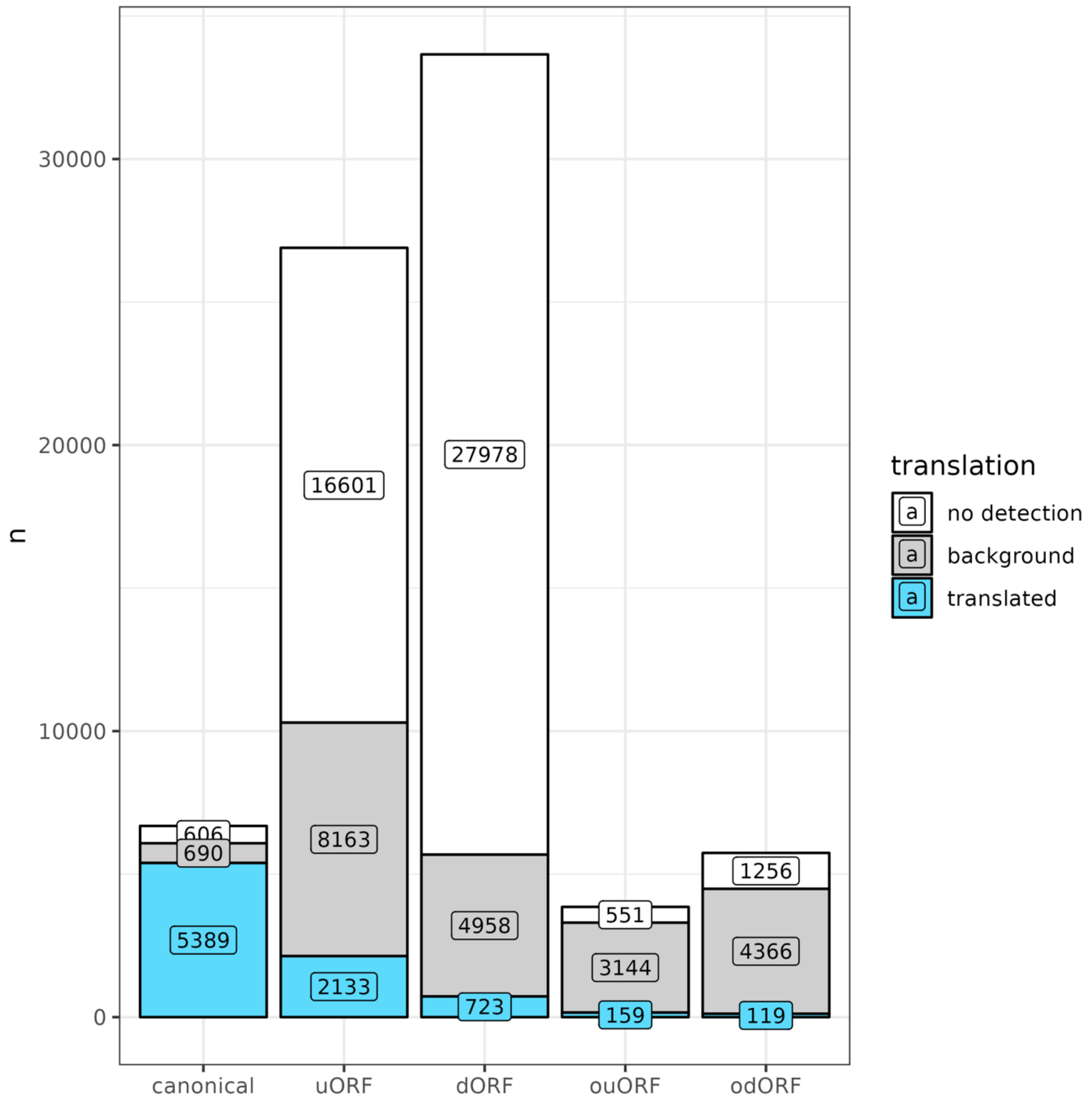


**Figure S2. ORFs in S. cerevisiae genes classified by translation status.** no detection: no Ribo-Seq reads mapping to the ORF; background: > 10 Ribo-Seq reads mapping to the ORF but not significant translation signature (ribORF score < 0.6); translated: > 10 Ribo-Seq mapped reads as well as translation signature (RibORF score ≥ 0.6).

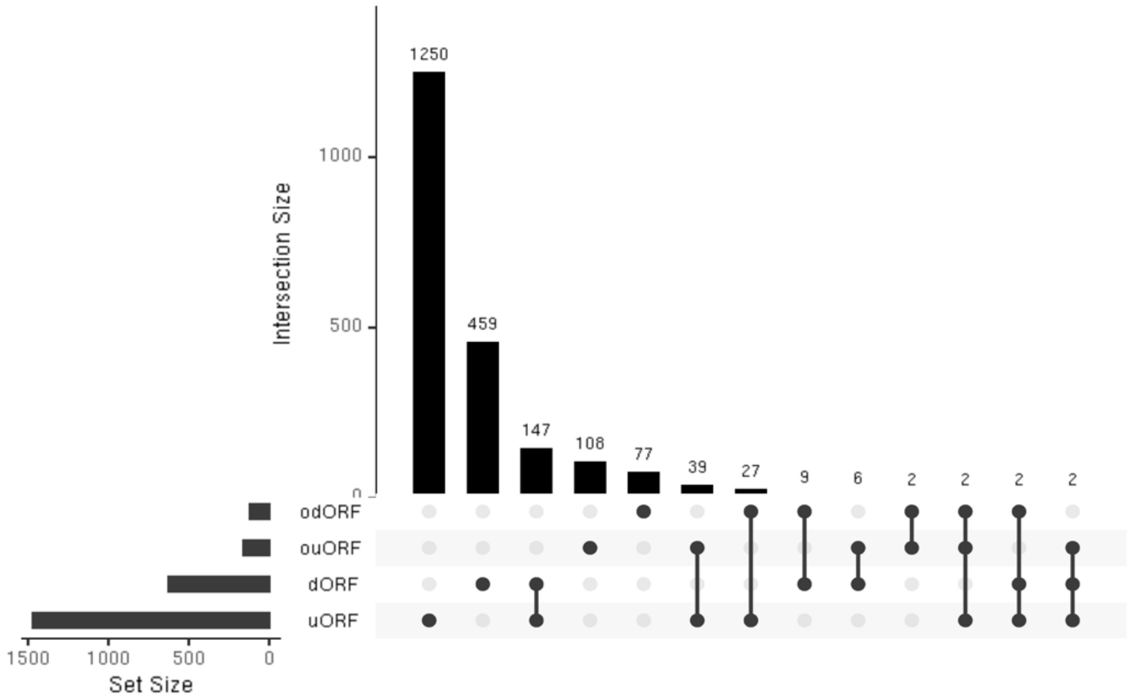


**Figure S3. *S. cerevisiae* genes containing translated ORFs in the UTRs**. The number of genes containing different only one type of ncORF or several types of ncORFs is shown. ncORF translation prediction was performed with RibORF v.2.0 (score cut-off > 0.6). Only genes with 5'UTR and 3'UTR identified using dRNAs were considered.

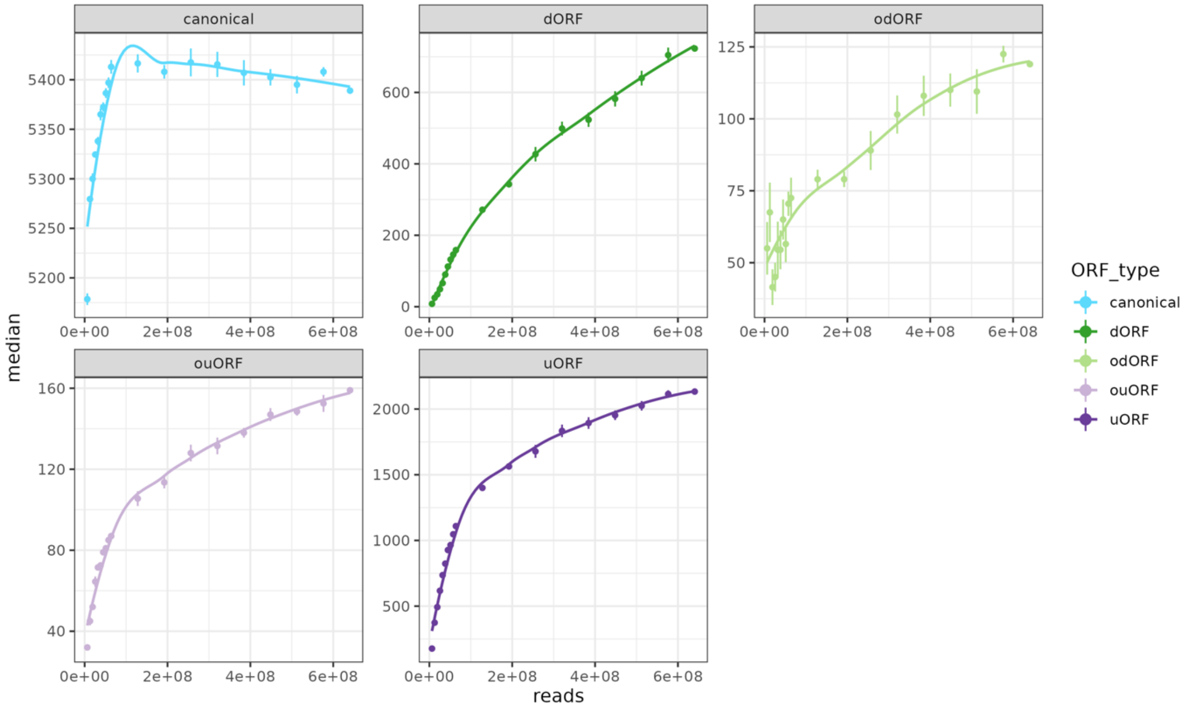


**Figure S4. Detail on the number of translation events detected with increasing number of Ribo-Seq reads.** Randomly selected increasing number of Ribo-Seq reads were taken and translated ORFs predicted. Canonical coding sequences show very rapid saturation, dORFs/odORFs are almost linear and uORFs/ouORFs show partial saturation.

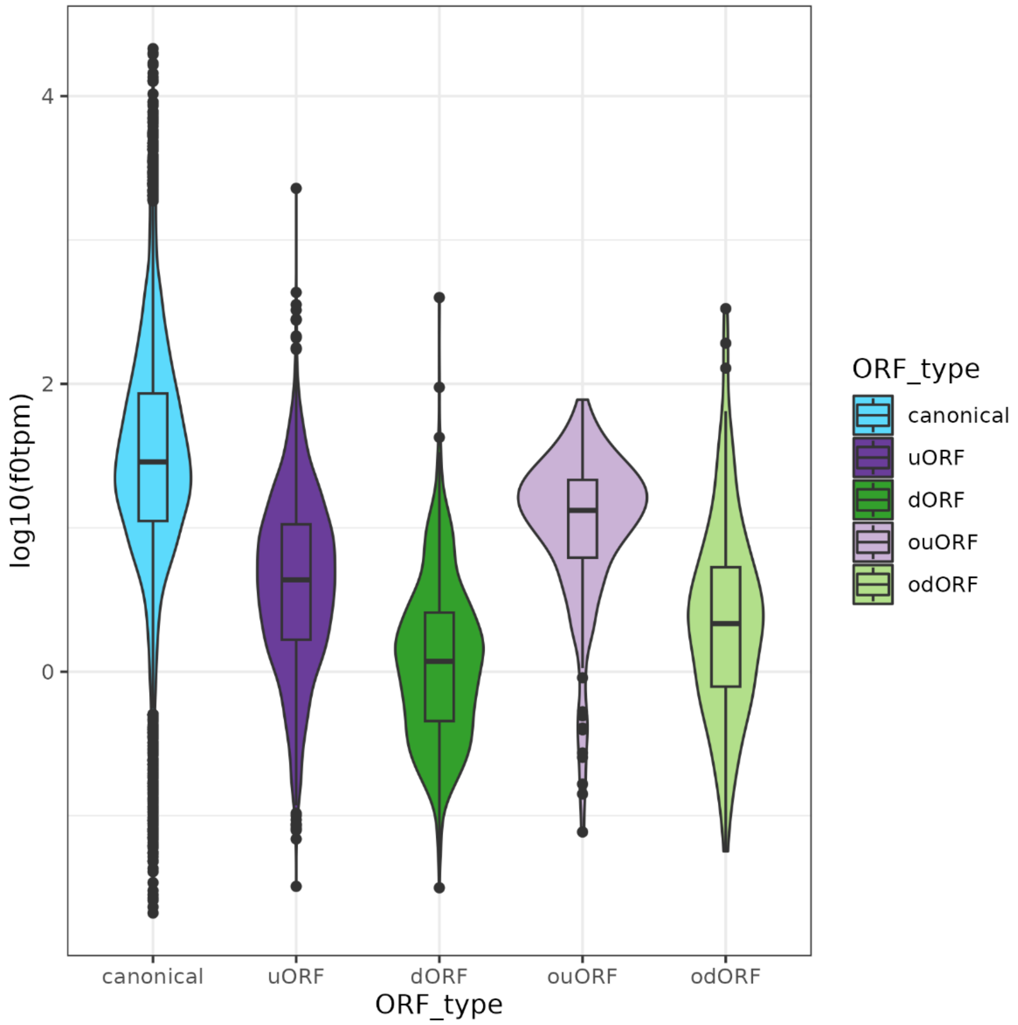


**Figure S5. Translation levels of different types of ORFs in *S. cerevisiae*.** Translation levels were calculated using TPM (transcripts per Million) with in-frame reads (f0tpm), which are those that correspond to the correct frame.

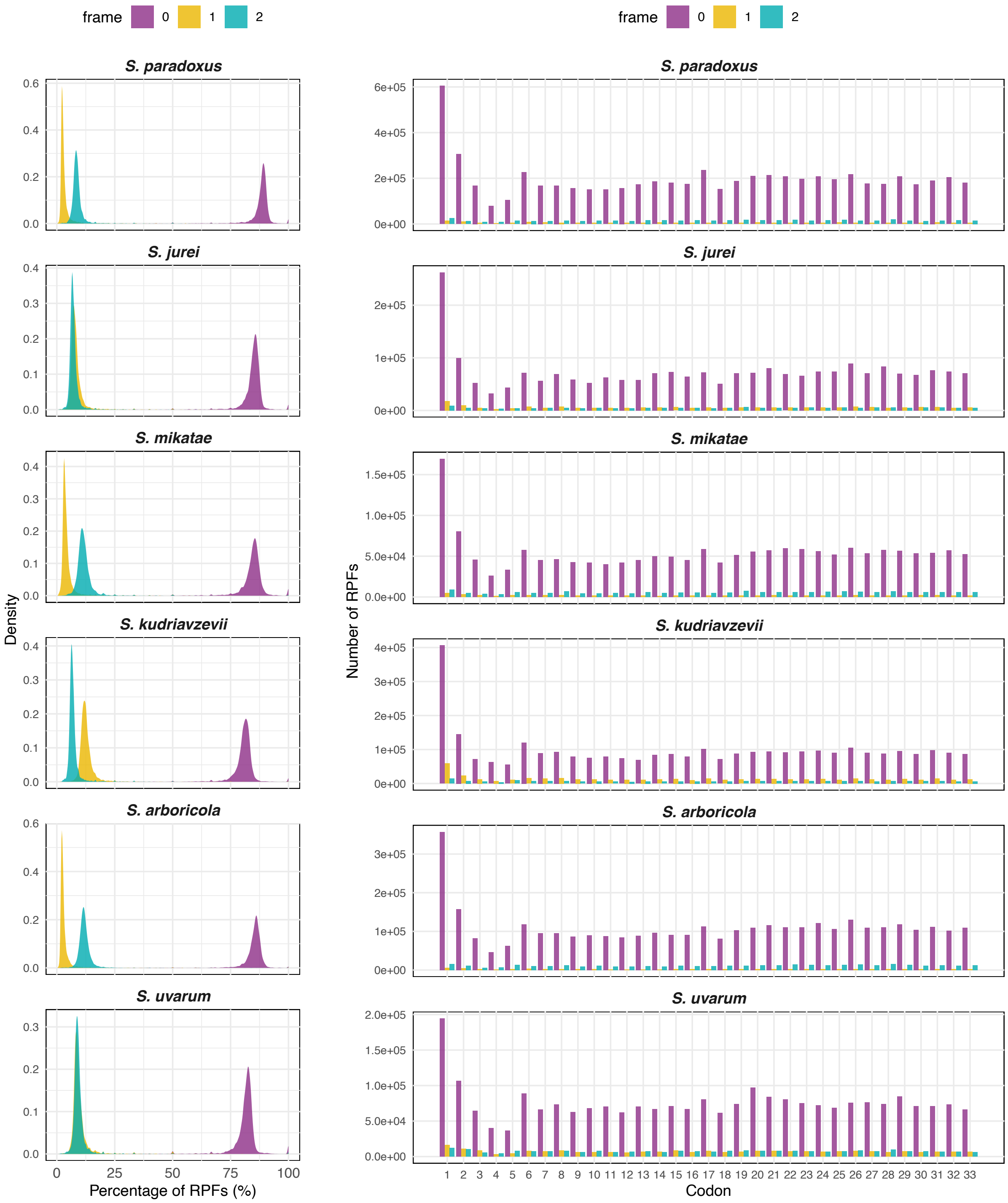


**Figure S6. Quality control of the Ribo-Seq data for six Saccharomyces species.** The data is for all the annotated CDS. The plots show that a very large fraction of the reads correspond to frame 0 (the correct frame).

| Gene Ontology term | Cluster frequency | Genome frequency | Corrected P-value | FDR | False Positives | Genes annotated to the term |
|---|---|---|---|---|---|---|
| transporter activity | 28 of 157 genes, 17.8% | 393 of 7166 genes, 5.5% | 3.61e-06 | 0.00% | 0.00 | YNR055C, YGL167C, YJR121W, YPL092W, YML116W, YLR378C, YGR065C, YBR291C, YMR026C, YPL221W, YMR034C, YER166W, YMR010W, YOR306C, YBL099W, YDR387C, YBR086C, YOR065W, YPR138C, YBR187W, YLR188W, YBR069C, YPR021C, YOL130W, YJR152W, YKR050W, YLR130C, YKL064W |
| transmembrane transporter activity | 25 of 157 genes, 15.9% | 351 of 7166 genes, 4.9% | 2.31e-05 | 0.00% | 0.00 | YOR306C, YBL099W, YDR387C, YML116W, YLR378C, YGR065C, YBR291C, YMR026C, YPL221W, YMR034C, YPL092W, YNR055C, YGL167C, YJR121W, YLR130C, YKL064W, YLR188W, YBR069C, YJR152W, YPR021C, YOL130W, YKR050W, YPR138C, YBR187W, YOR065W |
| phosphotransferase activity, alcohol group as acceptor | 16 of 157 genes, 10.2% | 183 of 7166 genes, 2.6% | 0.00036 | 0.00% | 0.00 | YHR079C, YNR012W, YDR173C, YJL128C, YPL153C, YOR061W, YDR247W, YOR231W, YER123W, YPL214C, YBR028C, YBL088C, YJL106W, YGL179C, YHR135C, YOL100W |
| protein kinase activity | 13 of 157 genes, 8.3% | 130 of 7166 genes, 1.8% | 0.00078 | 0.00% | 0.00 | YER123W, YOR231W, YDR247W, YOR061W, YJL128C, YPL153C, YHR079C, YOL100W, YHR135C, YGL179C, YJL106W, YBL088C, YBR028C |
| kinase activity | 16 of 157 genes, 10.2% | 199 of 7166 genes, 2.8% | 0.00108 | 0.00% | 0.00 | YER123W, YDR247W, YOR231W, YJL128C, YPL153C, YOR061W, YHR079C, YNR012W, YDR173C, YHR135C, YOL100W, YGL179C, YBR028C, YBL088C, YJL106W, YPL214C |
| transferase activity | 36 of 157 genes, 22.9% | 844 of 7166 genes, 11.8% | 0.00875 | 0.00% | 0.00 | YGL179C, YFL009W, YMR026C, YPL214C, YDR054C, YER091C, YBL088C, YBR028C, YOR231W, YJL139C, YKR004C, YER123W, YJL130C, YBR114W, YDR173C, YNR012W, YOR370C, YKR069W, YLR097C, YNR058W, YOL100W, YHR135C, YML106W, YML023C, YBL052C, YJL106W, YKL104C, YNL256W, YDR247W, YDL205C, YKL182W, YHR079C, YJL140W, YOR061W, YJL128C, YPL153C |

**Figure S7. Gene Ontology (GO) enrichment in genes containing uORFs translated in multiple species.** The Gene Ontology Term Finder at the Saccharomyces Genome Database (SGD) was used to retrieved enriched Functions and calculate the adjusted p-value.

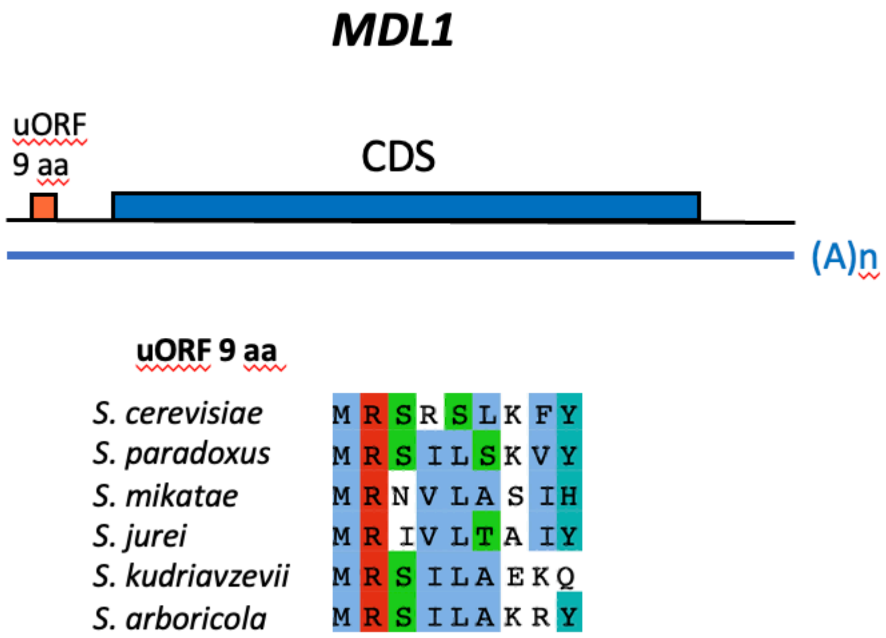


**Figure S8. Translated uORF in *MDL1*.** Microprotein translated from a uORF in MDL1 (YLR188W). The uORF encoding a 9 aa protein in *S. cerevisiae* was significantly translated in all species except *S. kudriavzevii* and *S. uvarum*. The latter has not been included in the alignment because the uORF was not conserved.

**uORF 27 aa in *PUS2***

```
S. cerevisiae    MKKTALKTSSVLRQIFTGQRIKRY--SYV
S. paradoxus     MKKAALKTGFALRHILTGQRIKRY--LYV
S. mikatae       MKNPVGTTSSVFRHIFRAQRIKKY--SYV
S. jurei         MKNAVVTTSSVFRHILRAQRIRTY--SYV
S. kudriavzevii  MKKTALDACPALRRSFAPQRIERH--LYVRMCMCIYVELL
S. arboricola    MKNTAIKAYSTLRQRFILQKSQKYKYSYVQAAIHGHGTDANISFY
S. uvarum        MKNFAVRACSSLPPRFAPQRIARY--SYV
```

**Figure S9. Alignment phylogenetically conserved uORF in *PUS2*.** The uORF encodes a protein of 27 aa in S. cerevisiae. Extensions can be observed in *S. kudriavzevii* and *S. arboricola*.

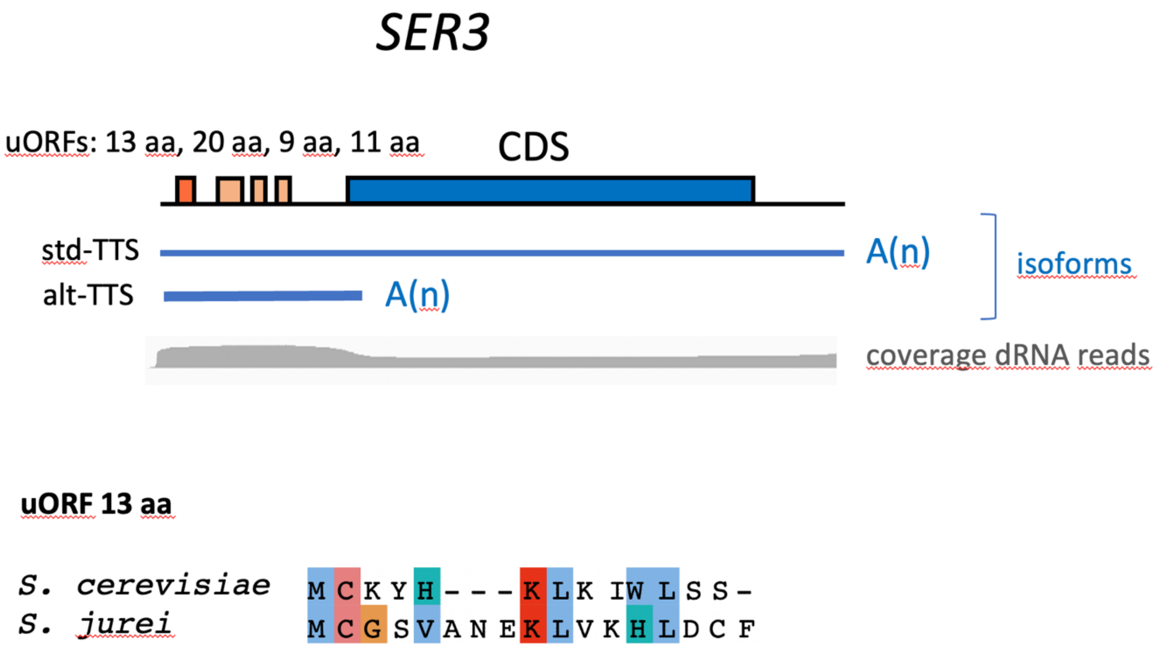


**Figure S10. Alternative isoform and translated uORFs in *SER3*.** We identified 4 translated uORFs in *S. cerevisiae*, one of which was also found translated in *S. jurei*. We identified an alt-TTS, generating an alternative isoform containing all four uORFs and associated with 2,256 reads, slightly more than the 1,878 reads assigned to the standard isoform (std-TTS).
